# Automated neuroimaging meta-analysis through LLM-guided selection improves recovery of published spatial patterns

**DOI:** 10.64898/2026.11.24.769029

**Authors:** Alejandro de la Vega, James D. Kent, Anthony Yang, Thomas E. Nichols, Jean-Baptiste Poline, Katherine L. Bottenhorn, Zijiao Chen, Russell A. Poldrack, Angela R. Laird

## Abstract

Systematic meta-analysis enables cumulative science, but expert-led, manual workflows do not scale with the growing literature. In neuroimaging, automated frameworks are widely used but typically sacrifice precision by using coarse article selection and pooling reported results, combining different tasks, populations or comparisons even when only a subset is relevant. Here we present AutoNIMA, a large language model (LLM)-guided framework that screens articles using expert-defined eligibility criteria and parses heterogeneous tables to extract brain stereotactic coordinates and select relevant analyses within each article. AutoNIMA produced maps more closely matching expert-curated references than a search-only baseline representing existing automated methods in 26 of 27 meta-analyses. Analysis-level selection accounted for most of the improvement. These findings demonstrate how LLM-guided curation can filter relevant analyses within the same paper and support scalable neuroimaging evidence synthesis.

## Introduction

The rapid expansion of the scientific literature makes scalable synthesis essential for researchers to integrate findings and stay abreast of new evidence ^1^. In neuroimaging, quantitative meta-analysis assesses spatial convergence in brain activation or structural differences across studies ^2^, helping researchers evaluate evidence beyond individual experiments. Despite the availability of integrated curation platforms ^3,4^, systematic efforts remain laborious. Estimates indicate biomedical syntheses require over a thousand research hours ^5^ and over a year on average to complete ^6^. As a result, automated methods have become widely used, supporting exploratory meta-analysis, large-scale cognitive mapping, and experimental findings ^7–11^.

Although automated meta-analytic methods have advanced from term-based text mining to predictive modelling and learned representations ^8,12–15^, two technical barriers limit their precision. First, semantic relevance of article text alone does not establish eligibility, which requires carefully interpreting participants, tasks, and experimental designs against explicit researcher-defined criteria ^16^. Second, neuroimaging articles report brain coordinates in tables containing multiple analyses with heterogeneous formatting, making automated extraction difficult. As a result, existing automated frameworks aggregate all analyses reported in the same article. This pooling combines distinct tasks, participant groups, or opposing contrasts and can include ineligible analyses, such as region-of-interest (ROI) results for whole-brain meta-analysis. Recent studies have demonstrated that large language models (LLMs) can screen biomedical studies against researcher-defined criteria with high sensitivity ^17–20^ —even matching or exceeding human screening sensitivity across Cochrane reviews ^21^. Their ability to interpret table structure alongside article context may also support extraction and selection of individual analyses. However, screening accuracy alone does not establish whether LLMs can extract and select individual neuroimaging analyses with expert-level accuracy or how automating these steps affects the fidelity of quantitative synthesis.

Here we present AutoNIMA, an LLM-guided framework that applies expert-defined eligibility criteria to neuroimaging articles and the individual analyses within them. AutoNIMA implements an end-to-end workflow encompassing literature search, article screening, coordinate extraction, analysis-level selection, and coordinate-based meta-analysis. Using NeuroMetaBench^22^, a companion benchmark, we assessed agreement with expert curation and compared automated maps with expert-reference maps for 27 meta-analyses. We isolated the contributions of article screening and analysis selection to determine which decisions improve spatial fidelity. By making this workflow broadly available, AutoNIMA accelerates targeted neuroimaging synthesis guided by expert-defined criteria.

## Results

### Overview of AutoNIMA and its evaluation

AutoNIMA uses expert-defined eligibility criteria to configure a sequential screening pipeline that yields a spatial consensus map for each target meta-analysis (Fig. 1). The workflow searches PubMed, screens abstracts, and then evaluates available full texts against more detailed article-level criteria. For eligible articles, heuristics locate candidate coordinate tables and LLMs separate their coordinates into individual analyses. Each analysis is then evaluated against analysis-level eligibility criteria using the surrounding article context. This distinction between article and analysis eligibility is central: a relevant research article may report several analyses, only some of which are eligible for a particular meta-analysis. Coordinates from selected analyses are submitted to coordinate-based meta-analysis to produce the final maps.

**Fig. 1.**
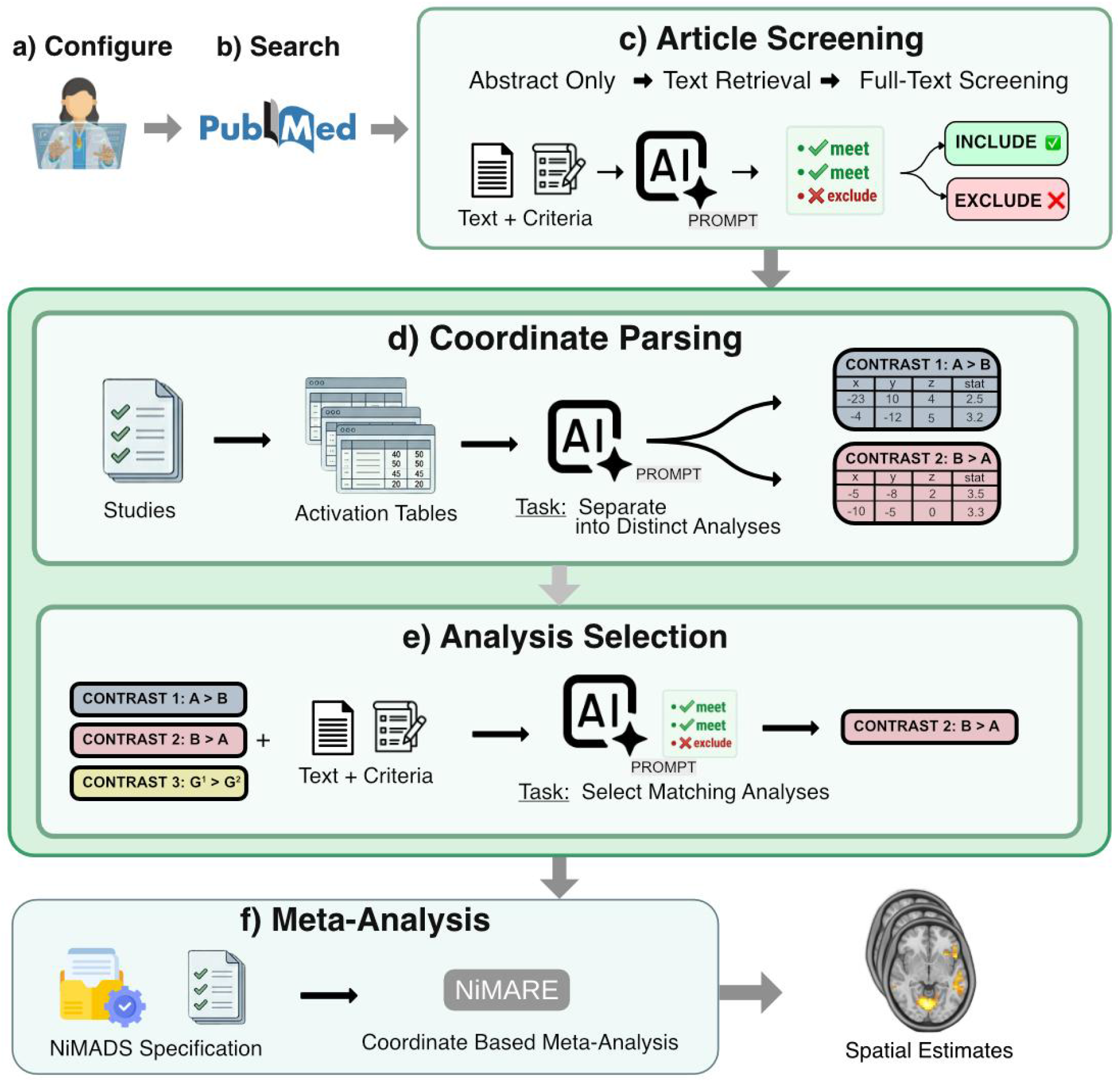
| AutoNIMA workflow from research question to meta-analytic map. a, Researchers specify article-level and analysis-level eligibility criteria for each meta-analysis. b, PubMed search retrieves candidate article records and abstracts. c, An LLM screens abstracts, then available full texts, returning eligibility decisions with supporting rationales and criterion-level assessments; articles proceed when all inclusion criteria are met and no exclusion criteria apply. d, Heuristics identify candidate coordinate tables, which LLMs parse into distinct analyses and coordinates. e, Each analysis is assessed against the eligibility criteria for each meta-analysis in the context of its research article and selected for the meta-analyses whose criteria it satisfies. f, Selected coordinates enter coordinate-based meta-analysis to generate a spatial convergence map for each meta-analysis.

We evaluated AutoNIMA using NeuroMetaBench, a companion benchmark drawn from nine quantitative review articles, described separately ^22^. Each review article in the benchmark reports multiple meta-analyses, each combining individual analyses from contributing research articles. Throughout, “expert” refers to the curated inclusion decisions, analyses and coordinates provided by NeuroMetaBench; “expert-reference maps” are maps reconstructed from those coordinates. The benchmark repository remained private during testing to minimize leakage of curated reference data, although the source publications were public. We manually assessed agreement with expert curation at each workflow stage and compared automated maps and expert-reference maps reconstructed using the same meta-analytic estimator. The fully automated, search-only baseline pooled all available parsed analyses from retrieved articles, without article screening or analysis-level selection, allowing us to evaluate the benefit of these stages.

For each review article, we encoded the expert eligibility criteria according to the reference publications and manually revised them to address ambiguities, omissions and transcription errors present in the article. Subsequent LLM-assisted refinement using source publications and benchmark error reports revealed additional omissions as well as inconsistently or underspecified criteria in the published studies. While benchmark-guided refinement yielded only a small improvement over manually tuned configurations (Fig. S1), LLM-assisted revisions largely increased alignment with the source publications. Thus, we report the configuration with the best screening performance against the expert inclusion lists, demonstrating the value of human-in-the-loop semi-automated configuration (see Supplementary Table S1 for full configuration matrix).

### Expert-defined criteria improve screening precision while retaining relevant studies

Screening and analysis-level selection progressively increased agreement with expert inclusion lists (Fig. 2a). Mean precision rose from 10.3% after PubMed search to 32.3% after full-text screening and 47.0% after retaining only articles with at least one selected analysis. Differences in the resulting search article pools from PubMed searches and the original reviews contributed to apparent false positives: in three reviews for which we were able to obtain the original candidate article lists, we used them as the screening pool and found that measured precision increased from 49.7% to 65.7% (Fig. S2; Methods). Mean recall decreased across screening stages but remained at 82.9% after analysis-level selection (Fig. 2b), excluding studies for which we were not able to retrieve the full text or extract coordinates from the main article body.

**Fig. 2.**
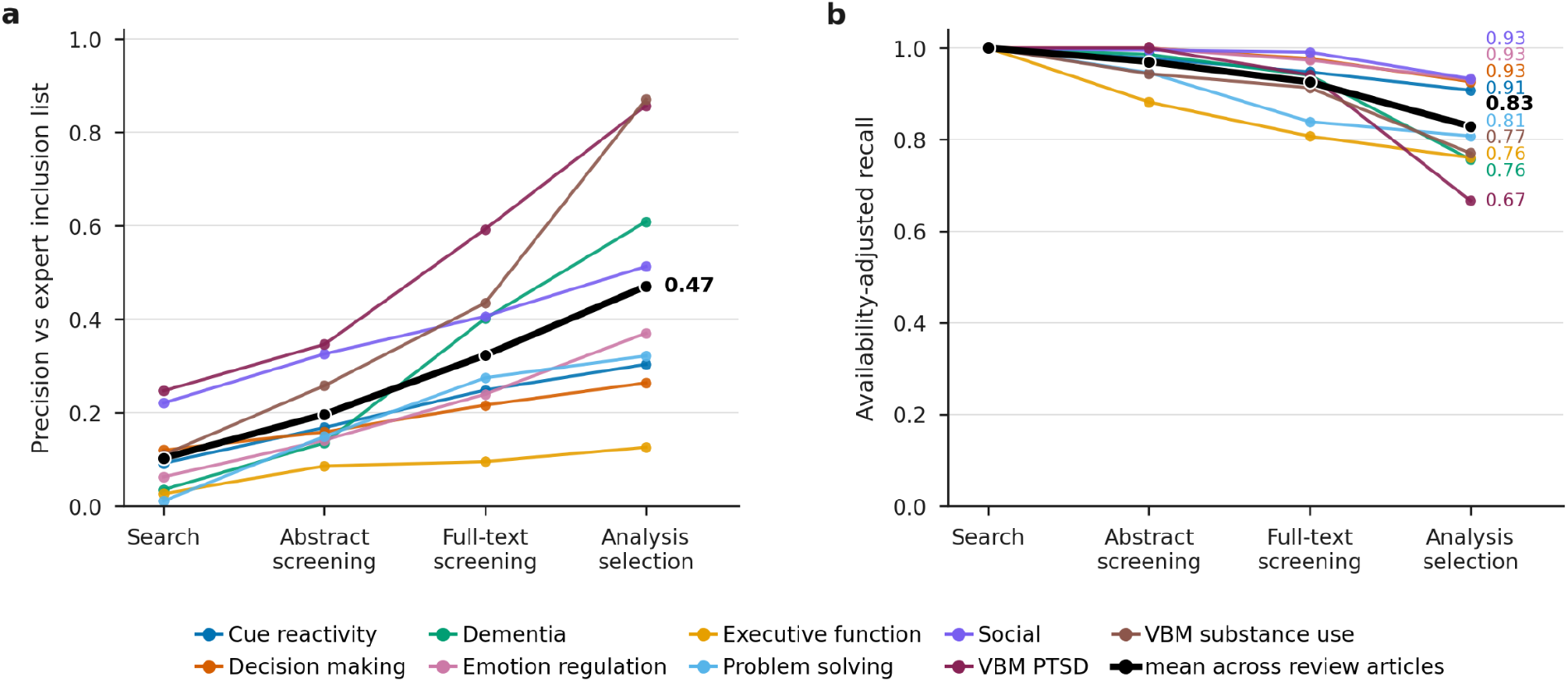
| Screening improves precision while retaining most available expert-included articles. **a**, Precision of included studies relative to the benchmark increased monotonically across stages of the pipeline. **b**, Availability-adjusted recall across the same stages decreased moderately across pipeline stages, with mean availability-adjusted recall of 82.9% at the final stage. Articles not retrieved by the search, lacking usable full text or yielding no parsed analyses were excluded from the denominator at the corresponding stage. Search-stage recall was therefore 100% by construction and does not indicate complete retrieval of the benchmark literature. In both panels, colored lines represent individual review articles (*n* = 9), and the mean metrics are represented in black.

### Automated coordinate parsing and analysis-level selection

We evaluated whether LLMs were able to group coordinates extracted from article tables into the same individual analyses as expert curators. Here, an analysis is an individual statistical result reported in a research article (e.g. a single contrast), represented by its coordinate set. Automatically parsed analysis records were matched to expert-curated records using coordinate and label similarity, with one-to-one assignment and predefined similarity thresholds (Methods). Among articles with extracted coordinates, we calculated the proportion of expert-curated analyses that we were able to match to an LLM parsed analysis. Across the nine review articles, this proportion averaged 91.5%, compared to a heuristic baseline of 33.2% when coordinates parsed from each table were grouped into a single analysis (Fig. 3a).

**Fig. 3.**
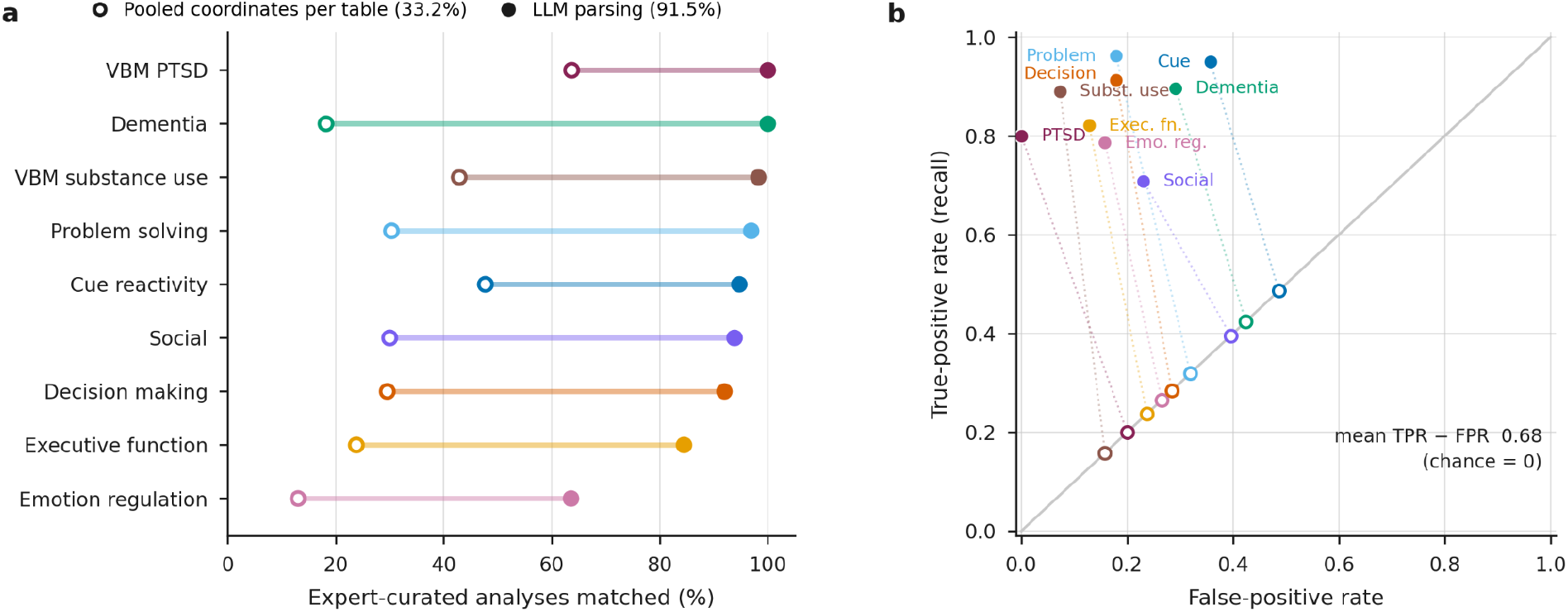
| LLMs parse coordinates into individual analyses and select them for meta-analyses in agreement with expert curation. a, Proportion of expert-curated individual analyses matched to automatically parsed analysis records. Matches used coordinate and label similarity with one-to-one assignment and predefined similarity threshold. Each marker represents one of the nine review articles. The matched proportion averaged 91.5% for LLM parsing (filled markers), compared with 33.2% when all coordinates parsed from each table were pooled into a single analysis (open markers). b, True-positive rate (recall) and false-positive rate for decisions to include an individual analysis in a meta-analysis. Each filled marker summarizes these decisions for one review article; a single analysis could be eligible for multiple meta-analyses. All nine review articles performed above the chance diagonal (mean TPR − FPR, 0.68). The diagonal indicates equal true-positive and false-positive rates under random selection. Open markers show the expected performance of uniformly selecting the same number of candidate assignments as AutoNIMA.

Next, for the automated analyses with an expert-curated match, we next evaluated whether AutoNIMA selected the same individual analyses for inclusion in each meta-analysis as experts. Each analysis–meta-analysis pair constituted a separate inclusion decision, because an analysis could contribute to more than one meta-analysis. Across 35 annotated meta-analyses from the nine review articles, mean assignment recall was 81.1% and precision was 53.9% (Fig. 3b; Methods). Assignment precision within each review article was 1.8–5.6 times that expected under random selection from the same candidate pool.

### Automated synthesis improves recovery of published spatial patterns

We next tested whether AutoNIMA’s end-to-end workflow translated improved evidence selection into more faithful recovery of published neuroimaging meta-analytic maps. To construct a representative baseline, we performed PubMed searches tailored to each meta-analysis in NeuroMetaBench, and pooled all resulting available analyses without subsequent article screening or analysis-level selection (‘search-only baseline’). Then, for both the AutoNIMA results and the baselines, we performed a coordinate-based meta-analysis (CBMA) using multilevel kernel density analysis (MKDA) ^23^, and computed the correspondence with expert-reference maps using squared voxelwise correlation between unthresholded maps (*r*²; Methods). The dementia review article was excluded because its reference data pooled coordinates across studies when they used the same dataset, producing analysis units that were not comparable to automatically extracted outputs.

Across 27 meta-analyses from eight review articles, AutoNIMA improved mean spatial correspondence by Δ*r*² = +0.124 (95% CI, +0.065 to +0.213, estimated by bootstrap resampling of review articles). Correspondence to expert-reference maps improved in 26 of 27 meta-analyses (Fig. 4b), mean *r*² was 0.564 for AutoNIMA and 0.440 for the search-only baseline. The higher *r*² was not due to simply including more analyses– AutoNIMA included fewer analyses than the search-only baseline in 19 of 27 meta-analyses.

**Fig. 4.**
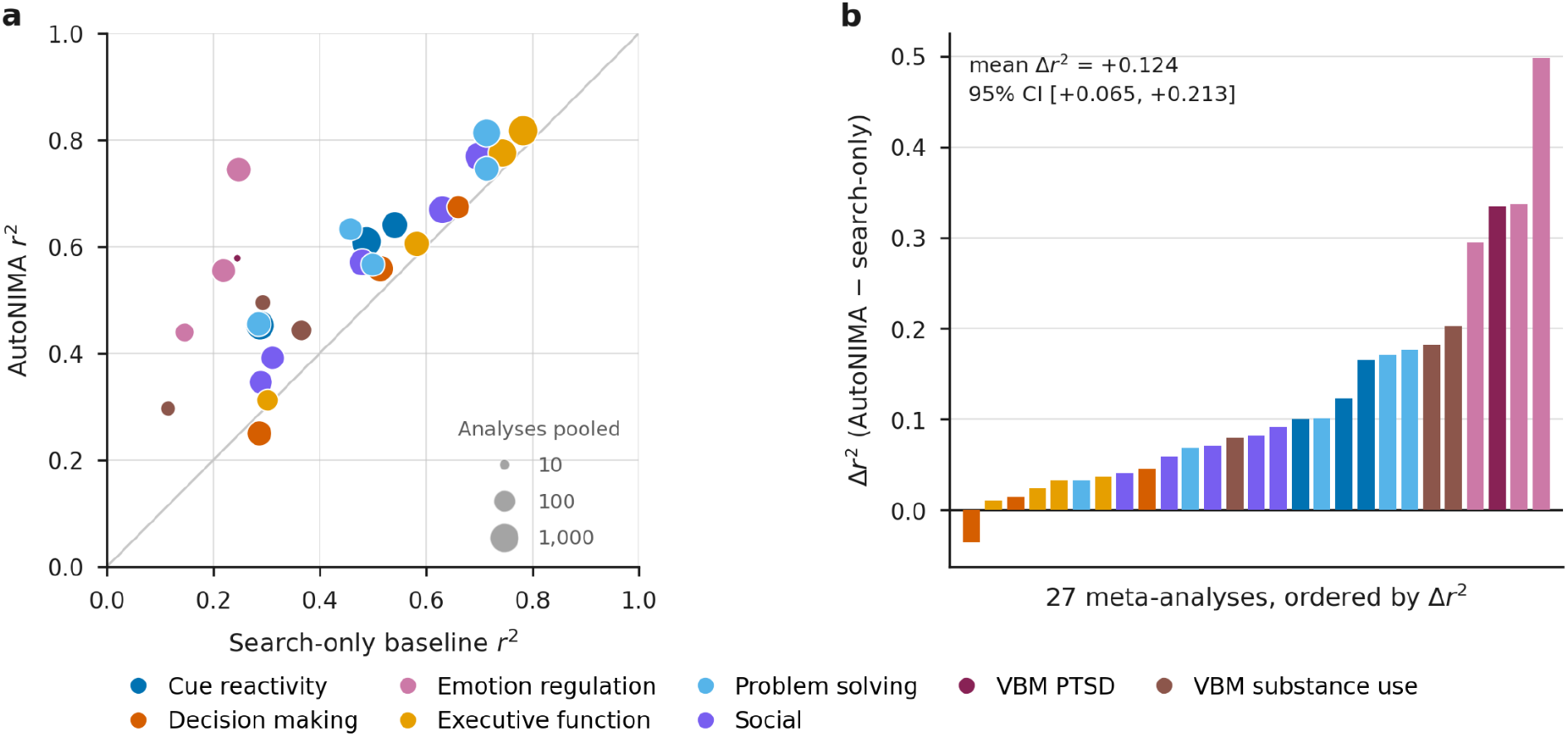
| AutoNIMA improves spatial correspondence with expert-reference maps. **a**, Correspondence of AutoNIMA and search-only baseline maps with expert-reference maps for 27 meta-analyses from eight review articles, measured as squared voxelwise correlation between unthresholded maps (*r*²). Search-only baselines pooled all available analyses from PubMed searches, targeted to individual meta-analyses where possible, without subsequent screening or analysis selection. Points above the identity line indicate AutoNIMA maps were more similar to the expert-reference maps than the search-only baseline. Marker area indicates the number of pooled analyses. b, Differences in spatial correspondence (Δ*r*²) between AutoNIMA and the search-only baseline for each of the 27 meta-analyses, ordered by magnitude. The annotation shows the mean difference and its 95% confidence interval, estimated by bootstrapping across the eight review articles. All maps were generated using multilevel kernel density analysis (MKDA). AutoNIMA achieved higher correspondence in 26 of 27 meta-analyses (mean *r*², 0.564 versus 0.440; mean Δ*r*², +0.124).

Improvements varied across review articles, with the best performance for the emotional regulation review and smaller gains for decision making and executive function. The meta-analysis of “decreasing emotional responses” in emotion regulation provided a clear example: the AutoNIMA map resembled the expert-reference map much more closely than the search-only baseline did (Δr² = +0.50; Dice 0.70 vs 0.42; Fig. 5). A single reappraisal study can report separate analyses involving increasing, decreasing or maintaining emotional responses. Article-level pooling combines coordinates across these analyses, whereas analysis-level selection identifies those relevant to each meta-analysis. Figure S3 shows the six meta-analyses with the largest gains and the six with the smallest gains relative to the search-only baseline.

**Fig. 5.**
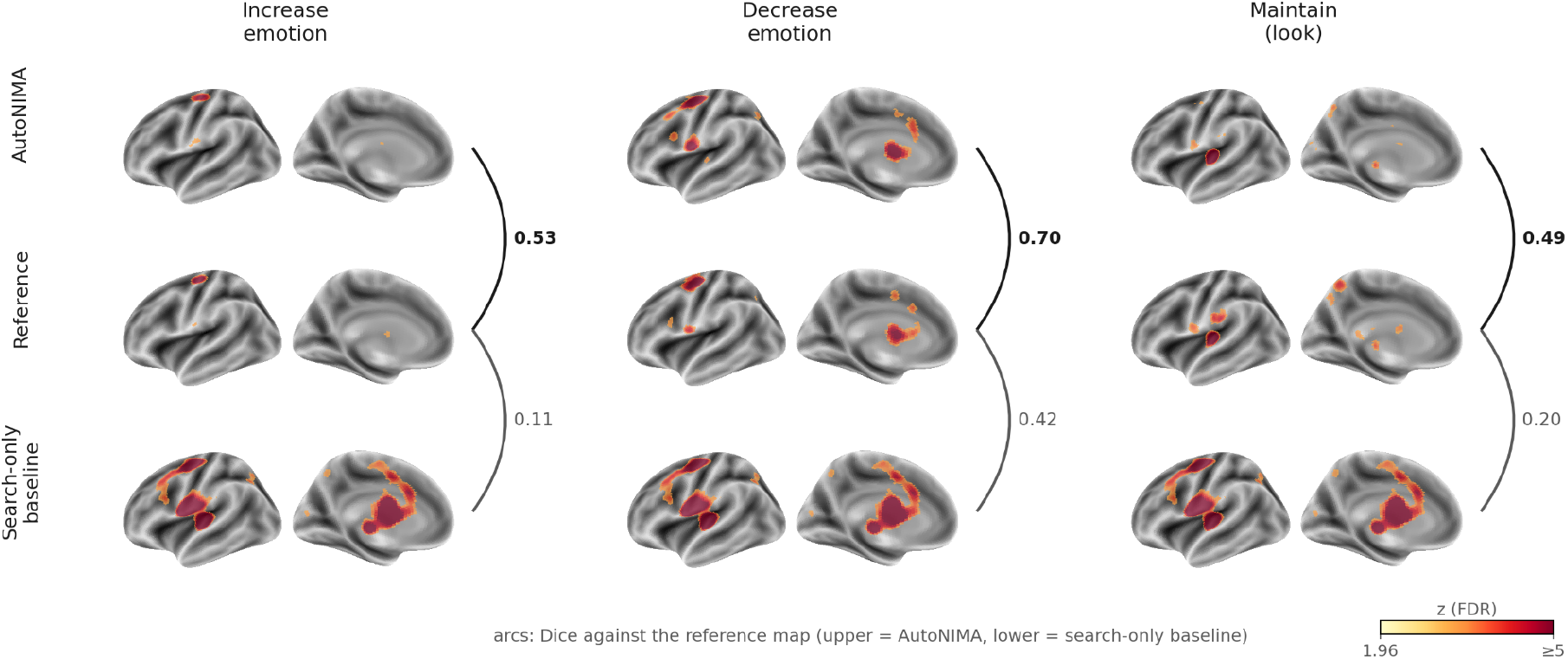
| Analysis-level selection distinguishes analyses relevant to different emotion-regulation meta-analyses. Rows show AutoNIMA, expert-reference and search-only baseline maps. Columns show the increase, decrease and maintain emotion-regulation meta-analyses, from left to right. Maps are displayed on inflated cortical surfaces at *z* > 1.96. A common emotion regulation search-only baseline is shown for all three meta-analyses; searches tailored to each meta-analysis produced similar maps. Dice coefficients quantify the spatial overlap of the search-only baseline and AutoNIMA maps with the expert reference.

To test whether our methodology depended on our specific pipeline, or if it instead could be orchestrated and executed by a more flexible domain-general AI harness (e.g. Claude Code, Codex), we reimplemented each workflow module—literature search, article screening, coordinate extraction, analysis selection, and meta-analysis—as a set of agent skills paired with equivalent criteria. Across three agent configurations evaluated on all nine review articles, screening agreement and spatial correspondence for available maps were broadly comparable across implementations (Fig. S6; Methods).

### Analysis selection drives spatial correspondence

Next, we assessed whether AutoNIMA’s improved spatial correspondence arose primarily from article screening or from parsing and selecting relevant analyses within articles. To test the contribution of article screening, we compared maps pooling all parsed analyses from screened articles without any analysis-level selection (“article selection only”) versus the same search-only baseline that pooled analyses from all studies. Screening added little spatial correspondence with the expert-reference maps (mean Δ*r*², +0.012; Fig. 6a), beyond searches that already targeted individual meta-analyses in 26 of 27 cases.

**Fig. 6.**
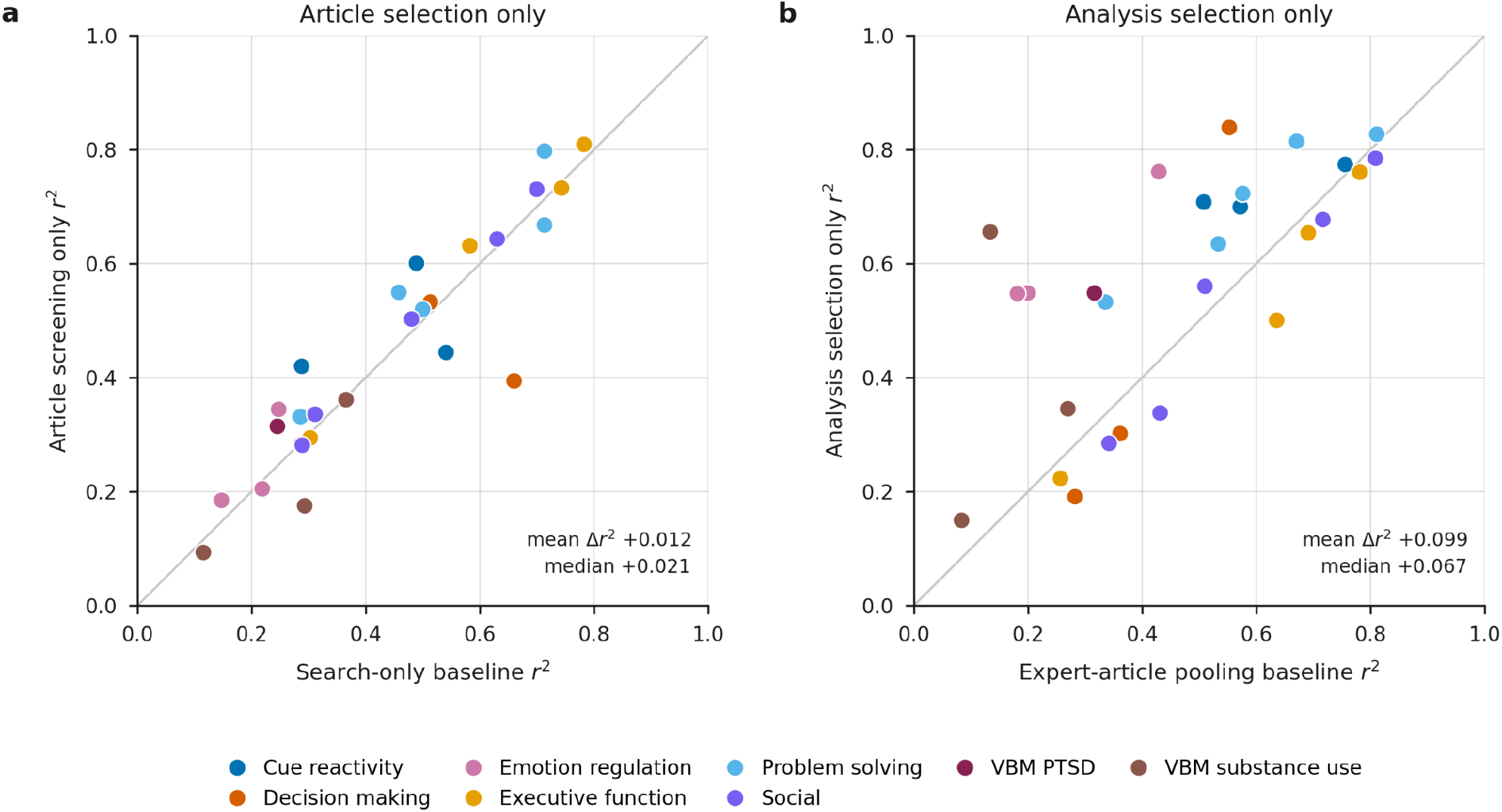
| Effects of article screening and analysis-level selection on spatial correspondence. **a,** Correspondence of article-screening-only maps and search-only baseline maps with expert-reference maps for 27 meta-analyses from eight review articles, measured as squared voxelwise correlation between unthresholded maps (*r*²). Both conditions pooled all available analyses from their respective article sets. **b**, Correspondence of analysis-selection-only maps and expert-article pooling baseline maps with expert-reference maps. Both conditions used the same expert-included articles from NeuroMetaBench; analysis selection retained results satisfying analysis-level eligibility criteria for each meta-analysis, whereas the baseline pooled all parsed analyses. In both panels, points above the identity line indicate improved correspondence after screening or selection. All maps were generated using multilevel kernel density analysis (MKDA).

To test whether analysis-level selection improved spatial correspondence after expert article screening, we applied AutoNIMA’s parsing and selection stages to a fixed pool of expert-screened articles from NeuroMetaBench (“analysis selection only”), and compared to a baseline that pooled all analyses from these expert-screened articles. Compared with the expert-article pooling baseline, analysis selection improved spatial correspondence in 17 of 27 meta-analyses (mean Δ*r*², +0.099; median, +0.067; Fig. 6b). Selected analyses also outperformed equally sized random subsets from the same pool in 24 of 27 meta-analyses (Fig. S4), suggesting that reducing the number of pooled analyses alone did not explain the improvement.

Finally, a separate paired comparison assessed the relative contributions of the two stages within the end-to-end workflow. Using the same initial article pool, analysis selection after screening contributed a mean Δ*r*² of +0.112, compared with +0.012 for screening alone, accounting for the total mean gain of +0.124 above the search-only baseline. Analysis selection contributed more than screening in 20 of 27 meta-analyses (mean difference in Δ*r*², +0.101; 95% CI, +0.038 to +0.190; Wilcoxon signed-rank *P* across review-article means = 0.016).

## Discussion

AutoNIMA leverages LLM agents to apply researcher-defined eligibility criteria within an integrated multi-stage workflow, enabling scalable automated and researcher-in-the-loop syntheses. Evaluating AutoNIMA against an expert-curated benchmark showed improved screening precision and identification of individual analyses, yielding greater spatial correspondence with expert-reference maps than a search-only baseline in nearly all evaluated meta-analyses. AutoNIMA’s analysis-level selection drove the improvement: article screening added little beyond targeted searches, whereas selecting relevant analyses considerably improved correspondence to the benchmark. Notably, applying AutoNIMA to analysis selection even improved on expert article screening, showing that identifying eligible papers is insufficient when those papers report multiple analyses addressing different questions. AutoNIMA applies LLM-driven eligibility assessment to all stages of a systematic neuroimaging meta-analysis, enabling targeted fully automated syntheses and accelerating human-in-the-loop curation.

The importance of analysis-level selection is highlighted in AutoNIMA’s performance on the emotion regulation review article. Across the three meta-analyses in this review, researchers sought to separate constituent components of emotion regulation. Included research articles often simultaneously reported analyses involving maintaining, increasing or decreasing emotional responses ^24^. The search-only baseline pooled these analyses into activation patterns that blurred key distinctions between conditions and, thus, distinct cognitive components of emotion regulation, even with expertly tailored keyword searches. A recent NeuroMetaBench evaluation identified the meta-analyses in this review article as challenging to reproduce for existing automated tools—including Neurosynth, NeuroQuery, and NeuroVLM— highlighting the limitations of methods pooling across reported analyses ^22^. By separating and selecting the relevant analyses, AutoNIMA recovered spatial patterns that more closely resembled the expert-reference maps.

AutoNIMA’s magnitude of improvement over search-only baselines nevertheless varied, with smaller gains in domains such as decision making and executive function. In some cases, this might be because broad research questions are adequately captured by the search-only baseline, leaving less opportunity for analysis-level selection to improve the resulting maps. Alternatively, smaller gains could reflect ambiguous construct definitions inherent to the research question, ^25^ imperfect expert-crafted criteria, or the limits of LLMs to approximate domain-expert judgment. Distinguishing these explanations would require several domain experts to carefully review the criteria and individual selection decisions for each review—an important next step that is beyond the scope of the present study.

Accurately specifying eligibility prompts was essential to AutoNIMA’s performance. Manual review of the transcribed criteria prompted revisions for five out of nine review articles, revealing transcription errors and ambiguous wording in published criteria (Fig. S1). Subsequent benchmark-guided refinement yielded only moderate improvement, but surfaced a different class of problem: flawed or inconsistently applied criteria in published meta-analyses. For instance, in the emotion regulation meta-analysis, LLMs faithfully applied the specified criteria to only include studies using static stimuli; however, the review’s authors ultimately included studies using movie stimuli. This pattern is consistent with findings demonstrating that LLM performance in scientific review depends not just on model capability but also accurate

specification, ^26^ and highlights the importance of human oversight and transparent reporting in AI syntheses ^27^. In neuroimaging, such best-practice AI guidelines are consistent with established recommendations for transparent reporting in systematic review ^16^, including clearly specifying participant characteristics, eligible analyses, and spatial search coverage.

Since seemingly minor specification errors can cascade to large downstream differences, iterative review of model judgment by researchers is essential. AutoNIMA facilitates this process by documenting screening decisions and providing interactive reports for post-hoc review of LLM decisions. Furthermore, we adapted the AutoNIMA workflow as a skills-based plug-in for Claude Code and OpenAI Codex and evaluated the implementation against the dedicated AutoNIMA harness (Fig. S6). This facilitates iterative LLM-guided criteria design, while simultaneously enforcing best-practice tracking of screening decisions and modifications to these criteria. Further community work should adapt best practice guidance to automated analysis-level curation, including procedures for evaluating extraction completeness, resolving ambiguous analysis descriptions and assessing how curation decisions affect resulting meta-analyses.

However, access to usable source data remains a major practical constraint on any automated synthesis approach. Only approximately one-third of NeuroMetaBench’s included articles could be retrieved automatically from open-access sources; additional coverage required manual retrieval or institutional access. In addition, only two-thirds of expert-included articles yielded extracted analyses in the main article body, with coordinates available only in supplementary files contributing to extraction failures. Depositing unthresholded statistical maps with clear analysis descriptions in repositories such as NeuroVault would reduce reliance on extracting coordinates from article texts ^28,29^ and facilitate more powerful methods such as image based meta-analysis ^30^.

Integration with existing meta-analysis platforms such as Neurosynth Compose ^3^ and NeuroVault could allow researchers to inspect, revise, and reuse automated selections across related questions. This approach aligns with proposals for living systematic reviews, which combine maintained searches with reusable, structured review data to reduce duplicated effort and incorporate new evidence ^31^. For neuroimaging, reusable selections could facilitate comparisons across populations, tasks, and contrast directions, and enable researchers to collaboratively test hypotheses in a shared environment. AutoNIMA provides a foundation for this approach by connecting researcher-defined criteria to analysis-level curation and quantitative synthesis.

## Methods

### AutoNIMA Pipeline

#### Configuration and execution

AutoNIMA implements an end-to-end workflow from literature search to coordinate-based neuroimaging meta-analysis. Each AutoNIMA project specifies a synthesis workflow in a YAML configuration containing a PubMed query, article-level inclusion and exclusion criteria, retrieval sources, parsing settings and analysis-level eligibility criteria for each meta-analysis. In this evaluation, each AutoNIMA project corresponded to one quantitative review article and specified one or more meta-analyses. Separate screening configurations allow broader criteria to be applied to abstracts and more detailed criteria to full texts. The Python pipeline executes the workflow sequentially, recording progress and outputs in a dedicated project directory. AutoNIMA caches interim outputs, enabling resumption of workflows and minimizing unnecessary computation and external API calls.

The four LLM-assisted stages—abstract screening, full-text screening, coordinate parsing, and analysis selection—access models through the OpenAI Python library. Model identifiers and API endpoints are configurable, allowing use of OpenAI-compatible providers. For this evaluation, all four stages used the general-purpose model gpt-5-mini-2025-08-07. Each stage used a task-specific prompt template combining general instructions with relevant user-defined configuration values, including project-specific eligibility criteria. Output formats were defined using Pydantic schemas compiled to JSON Schema, and model responses were validated against these specifications. Stage-specific outputs included parsed analyses and coordinates, eligibility decisions, criterion-level assessments, and supporting explanations.

#### Literature search

Candidate articles were identified through PubMed using the Entrez API and project-specific queries. Search queries, date restrictions and search dates were recorded in the project configurations and run records. Because the present evaluation focused on automated curation rather than search performance, we transcribed the published PubMed queries verbatim and refined them through expert review and LLM assistance to improve retrieval of benchmark included articles.

#### Article screening

The screening workflow proceeded from abstract screening to full-text retrieval and screening. Abstract screening provided a lower-cost initial filter for clearly ineligible articles, whereas full-text screening was performed using the more complete information available in the retrieved article. The same general screening procedure was used at both stages, with the information supplied to the model differing according to the stage. Project-specific screening criteria were specified in the pipeline configuration and incorporated into the screening prompts. In configurations used in this evaluation, abstract-screening criteria were intentionally more permissive to account for the incomplete information available in abstracts, whereas full-text screening used the complete set of project-specific inclusion and exclusion criteria.

At each screening stage, the model was instructed to evaluate the supplied article information against the configured criteria. For each inclusion criterion, the model reported whether the criterion was satisfied, and for each exclusion criterion, whether the criterion was triggered. The model was instructed to exclude an article if any required inclusion criterion was not satisfied or if at least one exclusion criterion was triggered. In addition to these criterion-level assessments, the model returned a Boolean eligibility decision and free-text reasoning explaining the decision. The Boolean decision determined whether the article proceeded to subsequent stages of the pipeline. Thus, articles were retained when all applicable inclusion criteria were satisfied and no exclusion criterion was triggered.

#### Full text retrieval

Full texts for articles that passed abstract screening were retrieved using sources specified in the project configuration. Supported sources include PubMed Central (PMC), publisher APIs, and user-provided article files. PubMed Central articles were retrieved using *pubget*, an open-source Python library for querying and fetching texts from PMC. Publisher retrieval is supported through text-mining APIs, including the Elsevier Full Text API and Springer Nature APIs; access to these services requires API registration using the applicable institutional or text-and-data-mining access rights. User-provided full-text HTML can also be supplied directly, including article HTML files obtained through institutional access. For the present evaluation, we supplemented automated retrieval with manually downloaded article files obtained using institutional access where possible. We were unable to include articles only available as PDF files or for which access could not be obtained.

Importantly, AutoNIMA distinguishes retrieval failures from eligibility decisions. Records for which no usable full text could be obtained are marked as *unavailable*, whereas articles for which the available text was incomplete were identified as *incomplete* by the LLM screener as part of the screening instructions. Neither condition was treated as an LLM screening error in the evaluation, because the failure concerned availability or quality of the retrieved article text rather than the model’*s* eligibility judgment.

#### Coordinate parsing into analyses

For articles retained after full-text screening, candidate coordinate tables were identified and extracted for further processing using deterministic heuristics. An LLM was then provided with each table, together with its caption and footnotes, and instructed to extract valid stereotactic coordinates and group them into the distinct statistical analyses represented in the table. The parsing prompt used a two-pass procedure: first identifying analysis-defining axes and result blocks, and then assigning each coordinate to the appropriate combination of these axes. Analysis boundaries were defined by explicit changes in contrast or comparison, contrast direction, activation or deactivation, participant group, treatment, task condition, session, time point, parametric effect, or other explicitly labeled analysis dimensions. The parser also handled analysis structure encoded in multi-level or repeated column headers, allowing a single table row to contribute coordinates to multiple analyses when the table reported multiple independent analysis blocks.

The parser was instructed not to treat descriptive anatomical subdivisions as separate statistical analyses unless the source explicitly identified them as distinct statistical maps. Similarly, separate whole-brain and region-of-interest result blocks were treated as distinct analyses when explicitly labeled. When no explicit analysis-defining distinction was present, the table was treated as a single analysis. The parser extracted coordinates only from explicitly identified spatial coordinate fields, required complete three-dimensional coordinate triples, and recorded the reported coordinate space when explicitly identified as MNI or Talairach. Where available, same-row statistical values were also retained with their reported statistic type.

### Analysis-level selection

Parsed analyses were evaluated against the analysis-level eligibility criteria for each meta-analysis using the full text of the corresponding research article. This evaluation was performed on an article-by-article basis: for each article, the LLM was provided with the full-text article together with all parsed analyses from that article and the complete set of analysis-level eligibility criteria for each meta-analysis. For each parsed analysis and each meta-analysis, the model evaluated whether the analysis satisfied the corresponding inclusion and exclusion criteria, producing a Boolean assignment for every analysis–meta-analysis pair. These decisions were represented as an analysis × meta-analysis Boolean inclusion matrix.

The criteria could include global criteria applicable to all meta-analyses within a project as well as criteria specific to individual meta-analyses. Global criteria were applied first as a hard gate: an analysis was excluded from a meta-analysis if a required global inclusion criterion was not satisfied or if a global exclusion criterion applied. Analyses passing the global criteria were then evaluated against the local inclusion and exclusion criteria for each meta-analysis. An analysis was assigned to a meta-analysis when all required criteria were satisfied and no applicable exclusion criterion was triggered. Because each analysis–meta-analysis pair was evaluated independently, a single analysis could be assigned to multiple meta-analyses when it satisfied the criteria for each. As with article-level screening, the model provided criterion-level assessments and a free-text rationale for each decision.

### Meta-analytic modeling

Coordinates from the selected analyses were submitted to coordinate-based meta-analysis separately for each synthesis. Selected analyses and their coordinates were represented in the NeuroImaging Meta-Analysis Data Structure (NiMADS), a standardized format for representing neuroimaging studies, analyses, and reported coordinates, enabling interoperability with the NiMARE library for neuroimaging quantitative meta-analysis. For each meta-analysis, the resulting coordinate set was used to generate a spatial meta-analytic map using MKDA. Automated, baseline and expert-reference maps were all reconstructed with the same MKDA procedure and settings, irrespective of the estimator used in the original publication. ^32^

NiMARE exposes a wide range of meta-analytic algorithms (KDA, MKDA, ALE) and multiple correction methods (FWE, FDR), which can be specified by users at runtime through the AutoNIMA interface. For this evaluation, maps were fitted with MKDA using NiMARE 0.21.0 via AutoNIMA 0.1.1, with NiMARE’*s* default configuration. Each coordinate was convolved with a sphere of 10 mm radius; where spheres overlapped, the maximum was retained; the resulting study-wise modelled activation maps were combined into an MKDA density statistic; and those summary statistics were converted to *p*-values against an approximate null distribution rather than by Monte Carlo resampling. Multiple comparisons were controlled by the Benjamini–Hochberg false discovery rate (FDR) procedure at *q* = 0.05. Maps were estimated using the MNI152 template, and a focus-counter diagnostic was run for each fit. For each meta-analysis, NiMARE produced an unthresholded *z* map and *p* maps and the FDR-corrected *z* and *p* maps and a cluster table; the two similarity metrics read the unthresholded and the corrected map respectively.

### Benchmark and Evaluation Procedure

We evaluated AutoNIMA using NeuroMetaBench, a companion benchmark drawn from nine quantitative review articles. For each review article, the benchmark provides article inclusion records and curated data identifying the analyses, coordinates, and analysis-to-meta-analysis assignments used in the reference synthesis. These records support evaluation of article screening, recovery of individual analyses, assignment of analyses to meta-analyses and spatial correspondence of the resulting maps. Benchmark construction is described in the companion report.

NeuroMetaBench contains annotations for 35 meta-analyses from nine quantitative review articles covering diverse topics. Three substance-use VBM meta-analyses—cannabis, opioids and stimulants—lacked expert-reference maps because the original review reported no significant clusters, leaving 32 with expert-reference maps. We excluded the emotion-regulation reappraisal meta-analysis because it is a superset of the decrease meta-analysis, retaining three emotion-regulation meta-analyses—increase, decrease and maintain—and 31 meta-analyses overall for spatial evaluation. The primary spatial evaluation additionally excluded four meta-analyses from the dementia review article, leaving 27 meta-analyses from eight review articles. The dementia reference records pooled coordinates across contributing studies, preventing consistent attribution to individual articles and comparison with article-level automated extraction. Dementia was retained in article screening and parsing evaluations and in the exploratory criteria-refinement analysis.

### Article screening

Article-level precision was defined as the proportion of retained articles present in the expert inclusion list. At the final analysis-selection stage, an article was considered retained if at least one of its analyses was selected for a meta-analysis.

Recall was adjusted for availability to distinguish screening decisions from failures of retrieval or extraction. The search-stage denominator comprised benchmark articles returned by the PubMed search, making recall at this stage 100% by construction. The same denominator was used for abstract screening. At full-text screening, benchmark articles lacking usable full text were removed from the denominator. At analysis selection, benchmark articles that reached coordinate extraction but yielded no parsed analyses were additionally removed. These availability-adjusted recall estimates describe retention conditional on the inputs available at each stage, rather than overall recovery of benchmark studies. Excluding articles yielding no parsed analyses may also exclude extraction failures, so these estimates should be interpreted alongside article coverage.

To examine the influence of the candidate article pool on measured screening performance, we repeated screening with the same eligibility criteria using candidate lists available from the original reviews in three review articles that made that list available: dementia, emotion regulation, and social cognition. We compared these results with screening of the articles returned by AutoNIMA’*s* PubMed searches. The emotion regulation list covered only part of the literature considered in the original review, limiting its use as a complete reconstruction of the original screening pool.

### Recovery of expert-curated analyses from table parsing

Automatically parsed analyses were matched to expert-curated records within the corresponding article using optimal one-to-one assignment with the Hungarian algorithm over pairwise similarities. Hungarian matching prevented a single automated analysis from matching multiple expert-curated records, and matched the two best matching automated-manual pairs. This was necessary in part because the expert-curated records contained only included analyses— that is, analyses that were not relevant to any of the meta-analyses were never entered by annotators. For the baseline, we pooled all coordinates parsed from each table into a single analysis and matched it to the expert-curated records using the same procedure. This kept the extracted coordinates fixed while removing the division of tables into individual analyses.

#### Pairwise similarity

Each candidate pair receives a combined score *s* = 0.70 · *s*_coord_ + 0.30 · *s*_label_. A greater weight was given to coordinates, as they are the most objective evidence of matching, whereas labels were annotator-assigned and not necessarily verbatim from article prose.

- ***s*_coord_, coordinate agreement.** Expert and automated coordinates are first paired by a second Hungarian assignment, this one minimising Euclidean distance in millimetres, which makes the score independent of the order coordinates appear in. Each paired distance *d* is mapped to a similarity by a piecewise-linear curve: 1.0 for *d* ≤ 1 mm, 0.9 for *d* ≤ 2 mm, then falling linearly to 0.6 at 4 mm, 0.2 at 8 mm and 0 at 12 mm and beyond. The mean of the paired similarities is multiplied by a coverage penalty, min(*n*_expert_, *n*_auto_) / max(*n*_expert_, *n*_auto_), so an automated analysis that recovers the right peaks but the wrong number of them is discounted in proportion. A bonus of 0.05 is added when the two coordinate sets are identical to one decimal place, and the result is clipped to [0, 1].
- ***s*_label_, analysis-name agreement.** The maximum ratio over four comparisons of the expert and automated analysis names — full against full, base against base, and each against the other’*s* base — where the base is the text before the first semicolon and both strings are case-folded and whitespace-normalised. Taking the maximum lets a fully-qualified label match a truncated one without penalty.

#### Acceptance

A pair counts as a recovered analysis at *s* ≥ 0.55. However, a coordinate-driven override additionally accepts a pair whose combined score is low but whose geometry is unambiguous, when any of the following holds: *s*_coord_ ≥ 0.90; *s*_coord_ ≥ 0.80 with the two coordinate sets of equal size and at least three coordinates; or equal-sized sets of at least three coordinates with ≥ 90% of pairs within 8 mm and a median paired distance ≤ 6 mm. The override exists because an automated analysis can reproduce an expert analysis exactly and still score poorly on the label, having inherited a table caption rather than an analysis name. Match at high confidence is defined as *s* ≥ 0.75, or a coordinate match satisfying the override criteria described above.

#### Coordinate space during matching

Coordinates in NeuroMetaBench were frequently converted to MNI space even if the original coordinates were reported as Talairach. To enable automated matching, we detected converted records based on if they reported decimal precision, and transformed the annotated coordinates back to Talairach space using NiMARE’*s* built in transformer. However, to ensure our assumption was correct, we scored the similarity of three variants of each such record — the coordinates as published, and both the Talairach-to-MNI and MNI-to-Talairach re-expressions — and kept the best-scoring one. The two converted variants are allowed to qualify as an exact set under an axis-wise tolerance of ± 1.0 mm, which the as-published variant is not. This transform is applied only for matching; it is independent of the space handling inside the meta-analysis (see *Meta-analytic modeling*).

### Analysis-level selection

To evaluate LLM performance in matching analyses to meta-analyses, we computed precision and recall of analysis-to-meta-analysis assignments relative to the expert-curated records, separately for each meta-analysis. Selection performance was evaluated at the level of analysis-to-meta-analysis assignments because a single analysis could satisfy the criteria for multiple meta-analyses.

#### Exhaustive matching strategy

The expert-curated records contained only analyses that were included in the reference meta-analysis; analyses that were excluded were not annotated, and their coordinates were therefore not available for comparison. Scoring only automated assignments that matched an expert annotation would consequently bias precision upward: although models could correctly match included analyses to the appropriate meta-analysis, they could also assign analyses that should have been excluded altogether. To conservatively account for these assignments, we used an exhaustive matching strategy. LLM assignments that did not match an expert annotation were counted as false positives only after all expert-annotated analyses from the corresponding article had been matched with high confidence to automated analyses. Because the benchmark did not annotate analyses that were present in an article but excluded from the reference meta-analysis, an unmatched automated analysis could not otherwise be assumed to represent a false-positive selection. Once all expert-annotated analyses had been matched, every remaining automated analysis entered the scoring universe for that meta-analysis: those the pipeline had assigned to the meta-analysis were counted as false positives, and those it had not were counted as true negatives.

Of the 11,138 analysis-to-meta-analysis assignments in the scoring universe, 5,787 (52.0%) were automated assignments with no matching expert annotation. Applying this strategy reduced pooled precision from 0.863 to 0.539 while leaving recall unchanged at 0.811.

#### Metrics

For each meta-analysis, precision, recall, and the corresponding ROC quantities were computed as

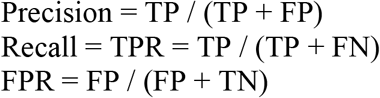

where TP, FP, FN and TN denote true-positive, false-positive, false-negative, and true-negative analysis-to-meta-analysis assignments, respectively. Pooled estimates were calculated by summing true positives, false positives, false negatives, and true negatives across review articles before computing the corresponding metrics. Receiver operating characteristic (ROC) displays for each review article plotted TPR against FPR.

#### Random-selection comparisons

Random-selection baselines were generated from the candidate assignment pool for each review article. For a fully labeled pool of *N* assignments, uniformly selecting *k* assignments yields an expected true-positive rate and false-positive rate of *k*/*N*, with expected precision equal to the prevalence of expert-positive assignments in the scoring universe. Precision enrichment was calculated relative to this prevalence. Random reference points were calculated analytically rather than by Monte Carlo sampling.

#### Coordinate-based meta-analysis and spatial correspondence

Meta-analytic maps were generated as described above (*Meta-analytic modeling*). The same estimation procedure and settings were used for the automated, baseline, and reference arms.

#### Spatial correspondence strategy

For each automated meta-analysis, spatial correspondence was evaluated by comparing its resulting statistical map with an expert-reference map reconstructed from the expert-curated coordinate records. Expert-reference maps were reconstructed using the same MKDA procedure, irrespective of the method used in the original publication (e.g., ALE). The comparisons therefore assess correspondence with reconstructions from published curation records rather than exact reproduction of the statistical analyses reported in the source publications.

Correspondence was also evaluated relative to a search-only baseline. The baseline included every analysis parsed from every article returned by the literature search for the meta-analysis, without LLM screening or analysis-level selection, and applied the identical meta-analytic procedure. For meta-analyses with a dedicated search, that targeted search defined the baseline; for meta-analyses that were not separable search topics, such as effect direction within a single paradigm, the broad search for the review article was used.

The effect of automated selection was expressed as the difference in correspondence between the automated and baseline arms:

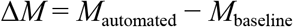

where *M* denotes the spatial correspondence metric (*r*² or Dice).

#### Spatial correspondence metrics

The primary measure was the squared Pearson correlation (*r*²) between unthresholded statistical maps. All Pearson correlations between unthresholded maps were calculated within an MNI152 brain mask at 2 mm resolution comprising 228,483 voxels.

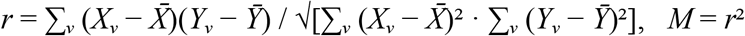

Dice overlap was used as a secondary measure for thresholded maps and illustrative map comparisons. Maps were thresholded at *z* > 1.96 using the FDR-adjusted *z* images, and Dice overlap was calculated as

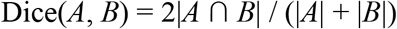

where *A* and *B* denote the sets of suprathreshold voxels.

#### Random-subset comparison

For each meta-analysis, we drew 500 random subsets from the parsed analyses in the expert-included article pool used for the analysis-selection-only comparison. Each subset contained the same number of distinct analyses as the observed selected set. MKDA was repeated using the same settings and expert-reference map, holding the number of pooled analyses constant while varying their identities.

Observed *r*² was compared with the mean and 5th–95th percentile range of the random-subset distribution. The one-sided empirical *P* value was calculated as (*b* + 1)/501, where *b* was the number of random subsets with correspondence at least as high as the observed result. We report unadjusted empirical *P* values and Benjamini–Hochberg-adjusted *q* values across the 27 meta-analyses. Percentile ranges describe variation among random subsets, not confidence intervals for the observed estimate.

#### Criteria refinement and benchmark feedback

Initial criteria for each review article were drafted to follow the eligibility descriptions in the source publications. We registered configurations in three tiers: initial, manually revised and benchmark-tuned. For each review article, the registry recorded the configuration versions, associated runs, and the reason for selecting the configuration used in the principal evaluation (Supplementary Table S1).

Manual inspection identified two errors in the initial eligibility specifications. The dementia specification omitted an eligibility criterion, and the cue-reactivity specification incorrectly introduced a restriction to healthy adults. Review of the corresponding model decisions confirmed that the models had applied the supplied criteria correctly. The manual revisions addressed these specification errors before benchmark-guided refinement. Subsequent revisions used LLM assistance and benchmark feedback, but as noted in Fig. S1, subsequent automated refinement had little effect on final map correspondence.

### Statistical inference and uncertainty estimation

Statistical inference for the end-to-end improvement in spatial correspondence and the paired comparison of workflow-stage contributions treated the review article as the independent unit. Meta-analyses within a review article share article pools, screening decisions and reference records and therefore are not independent observations.

#### AutoNIMA versus the search-only baseline

End-to-end change in map correspondence (Δ*r*²) was summarized across 27 evaluated meta-analyses from eight review articles. The 95% confidence interval for the mean change was estimated using a percentile bootstrap resampling review articles with 20,000 resamples (random seed = 0). Review articles were sampled with replacement, retaining all meta-analyses within each selected review article, and the mean change across meta-analyses was calculated for each resample.

#### Component comparisons

Article screening and analysis-level selection were evaluated against the search-only baseline and the expert-article pooling baseline, respectively (Fig. 6). The screening comparison used articles retrieved by PubMed searches, whereas the analysis-selection comparison used expert-included articles from NeuroMetaBench to assess the benefit of analysis selection after article eligibility had been resolved. Changes in spatial correspondence were summarized descriptively; these effects were not directly compared because they were measured in different article pools.

#### Paired decomposition

In a separate analysis, the search-only, screening-only and full-pipeline arms used the same initial article pool. The contribution of screening was defined as screening-only minus search-only correspondence, and that of analysis selection as full-pipeline minus screening-only correspondence. These sequential contributions summed to the end-to-end change for each meta-analysis. Their difference was evaluated using bootstrap resampling of review articles as described above and a Wilcoxon signed-rank test across the eight mean differences calculated within review articles. The expert-included article arm in Fig. 6b was excluded from this decomposition.

### Skills-based agent comparison

We reimplemented each AutoNIMA module—literature search, article screening, coordinate extraction, analysis selection and meta-analysis—as skills executed by a general-purpose orchestrator agent. All components of AutoNIMA’s execution harness, including its PubMed E-utilities client, full-text normalization scripts and NiMARE integration, were made available to the orchestrator agents. A decision-ledger script was added to support orchestration across modules and agents; it recorded all judgments, validated response structure and criterion identifiers, and verified quotations against source text. For stages requiring LLM judgment, the orchestrator launched judge agents without tool access, which returned structured responses.

Three orchestrator–judge configurations were evaluated on all nine review articles: Sol–Luna 5.6 and Astra–Astra in Codex, and Claude Opus 5.5–Claude Haiku in Claude Code. All configurations used the same eligibility criteria, search queries, date limits, map-generation settings, inclusion rules and evaluation procedures as the dedicated AutoNIMA harness (Fig. S6). The skills-based comparison retained the original evaluation set, including the emotion-regulation reappraisal meta-analysis, which was excluded from the primary spatial evaluation because it overlaps with decrease. Expert-reference files were kept outside the agent workspace.

Execution behavior was recorded for all configurations, whereas token usage was recorded only for Opus–Haiku. After optimization, recorded token usage for Opus–Haiku was moderately higher than for the dedicated AutoNIMA harness. In practice, we observed that runs frequently exhausted coding-agent subscription usage allowances, extending completion time. Opus–Haiku required one to four operator decisions per review article concerning unverified quotations, retry limits or meta-analyses with few contributing studies; Codex configurations rarely paused.

### Model-use costs

Model-use costs were estimated from recorded input, cached-input, and output token consumption using per-stage token accounting (Fig. S5). Abstract screening and full-text screening usage were measured over 708 and 1,211 model calls, respectively, across two review articles each; coordinate parsing and analysis selection were measured over 1,299 and 1,091 calls, respectively, across three review articles each. Token counts were converted to US dollar costs using the published gpt-5-mini-2025-08-07 per-token prices as of 1 October 2026, applying the respective rates for uncached input, cached input and output tokens.

For the skills-based implementation, costs were estimated only for Opus–Haiku, for which token usage was recorded. Input, cached-input and output rates were US$1, US$0.10 and US$5 per million tokens for Claude Haiku, and US$4, US$0.20 and US$20 for Claude Opus 5.5, respectively.

## Data availability

Project configurations, pipeline outputs (search results, screening decisions, parsed coordinates and analysis assignments) and evaluation tables are available at https://github.com/neurostuff/autonima-paper/. This repository and the generated maps are archived on Zenodo ^33^. Expert-curated reference data are available from NeuroMetaBench v1.0^34^.

## Code availability

AutoNIMA is available under the MIT License at https://github.com/neurostuff/autonima. Code used to construct the benchmark is available at https://github.com/neurostuff/neurometabench/.

## Use of generative AI

The use of LLMs for screening, coordinate parsing, and analysis selection constituted the method evaluated in this study and is described above. Separately, the authors used OpenAI Codex and Anthropic Claude to assist with software development of the main AutoNIMA package. The authors also used Anthropic Claude to generate evaluation scripts. During the preparation of this manuscript, the authors utilized OpenAI GPT-6 Astra to assist with editing and revising text for clarity, readability, and flow. In all instances, the AI-generated text was critically reviewed, edited, and revised by the authors to ensure accuracy and originality. The authors take full responsibility for all content presented in this manuscript, including any portions assisted by AI.

## Author contributions

A.V. conceived the study, developed the methodology, led software development and formal analysis, supervised the research team and wrote the original draft. J.D.K. helped shape the study aims, developed core software tools and reviewed and edited the manuscript. A.Y. developed supporting software and analysis scripts, helped revise the evaluation configurations and supervised individual subprojects. Z.C. helped run the computational analyses. T.E.N., J.B.P., K.L.B., R.A.P. and A.R.L. helped shape the study aims and reviewed and edited the manuscript.

