## Supplementary Information for "Automated neuroimaging meta-analysis through LLM-guided selection improves recovery of published spatial patterns"

### Supplementary Table S1 | Details of criteria revisions across initial, manually revised, and benchmark-tuned configurations.

| Review article | Configuration | Run | Mean $r^2$ | Criteria changed | Revision type | Revised by | Reported |
|---|---|---|---|---|---|---|---|
| Cue reactivity | Initial | v1 | 0.336 | | | | |
| Cue reactivity | Manually revised | v4 | 0.552 | 5 | Mis-transcribed criterion corrected | Manual (source publication) | |
| Cue reactivity | Benchmark-tuned | v6 | 0.568 | 18 | Ambiguous criterion clarified | LLM-assisted (source publication, error reports) | Yes: Highest screening F1 and recall |
| Decision making | Initial | v1 | 0.489 | | | | |
| Decision making | Manually revised | v2 | 0.497 | 4 | Ambiguous criterion clarified | Manual (source publication) | |
| Decision making | Benchmark-tuned | v3 | 0.495 | 10 | Omitted criterion added | LLM-assisted (source publication, error reports) | Yes: Highest screening F1 |
| Dementia | Initial | v1 | 0.06 | | | | |
| Dementia | Manually revised | v3 | 0.292 | 18 | Omitted criterion added | Manual (error reports) | Yes: Highest screening F1 |
| Dementia | Benchmark-tuned | v4 | 0.34 | 6 | Criterion contradicted by the source's included studies | LLM-assisted (source publication, error reports) | |

| Review article | Configuration | Run | Mean $r^2$ | Criteria changed | Revision type | Revised by | Reported |
|---|---|---|---|---|---|---|---|
| Emotion regulation | Initial | v2 | 0.545 | | | | |
| Emotion regulation | Manually revised | — | — | — | Not revised before tuning (same as initial) | | |
| Emotion regulation | Benchmark-tuned | v4 | 0.58 | 23 | Criterion contradicted by the source's included studies (revised from initial) | Manual (error reports) | Yes: Highest screening recall |
| Executive function | Initial | v1 | 0.609 | | | | |
| Executive function | Manually revised | — | — | — | Not revised before tuning (same as initial) | | |
| Executive function | Benchmark-tuned | v3 | 0.628 | 28 | No direct source basis (revised from initial) | LLM-assisted (source publication, error reports) | Yes: Highest screening recall |
| Problem solving | Initial | v1 | 0.59 | | | | |
| Problem solving | Manually revised | — | — | — | Not revised before tuning (same as initial) | | |
| Problem solving | Benchmark-tuned | v2 | 0.643 | 17 | Omitted criterion added (revised from initial) | LLM-assisted (source publication, error reports) | Yes: Tied screening F1; corrected analysis-level criteria |
| Social cognition | Initial | v2 | 0.501 | | | | |
| Social cognition | Manually revised | v3 | 0.55 | 0 | Search query changed (criteria unchanged) | Manual (source publication) | Yes: First configuration run on full texts from its own search |
| Social cognition | Benchmark-tuned | v4 | 0.548 | 3 | No direct source basis | LLM-assisted (source publication, error reports) | |

| Review article | Configuration | Run | Mean $r^2$ | Criteria changed | Revision type | Revised by | Reported |
| --- | --- | --- | --- | --- | --- | --- | --- |
| VBM of PTSD | Initial | v1 | 0.578 | | | | Yes |
| VBM of PTSD | Manually revised | — | — | — | Not revised before tuning (same as initial) | | |
| VBM of PTSD | Benchmark-tuned | v2 | 0.578 | 3 | Omitted criterion added (revised from initial) | LLM-assisted (source publication, error reports) | |
| VBM of SU | Initial | v1 | 0.42 | | | | |
| VBM of SU | Manually revised | v2 | 0.412 | 7 | Mis-transcribed criterion corrected | Manual (source publication) | Yes: Higher screening recall than initial |
| VBM of SU | Benchmark-tuned | v3 | 0.375 | 9 | No direct source basis | LLM-assisted (source publication, error reports) | |

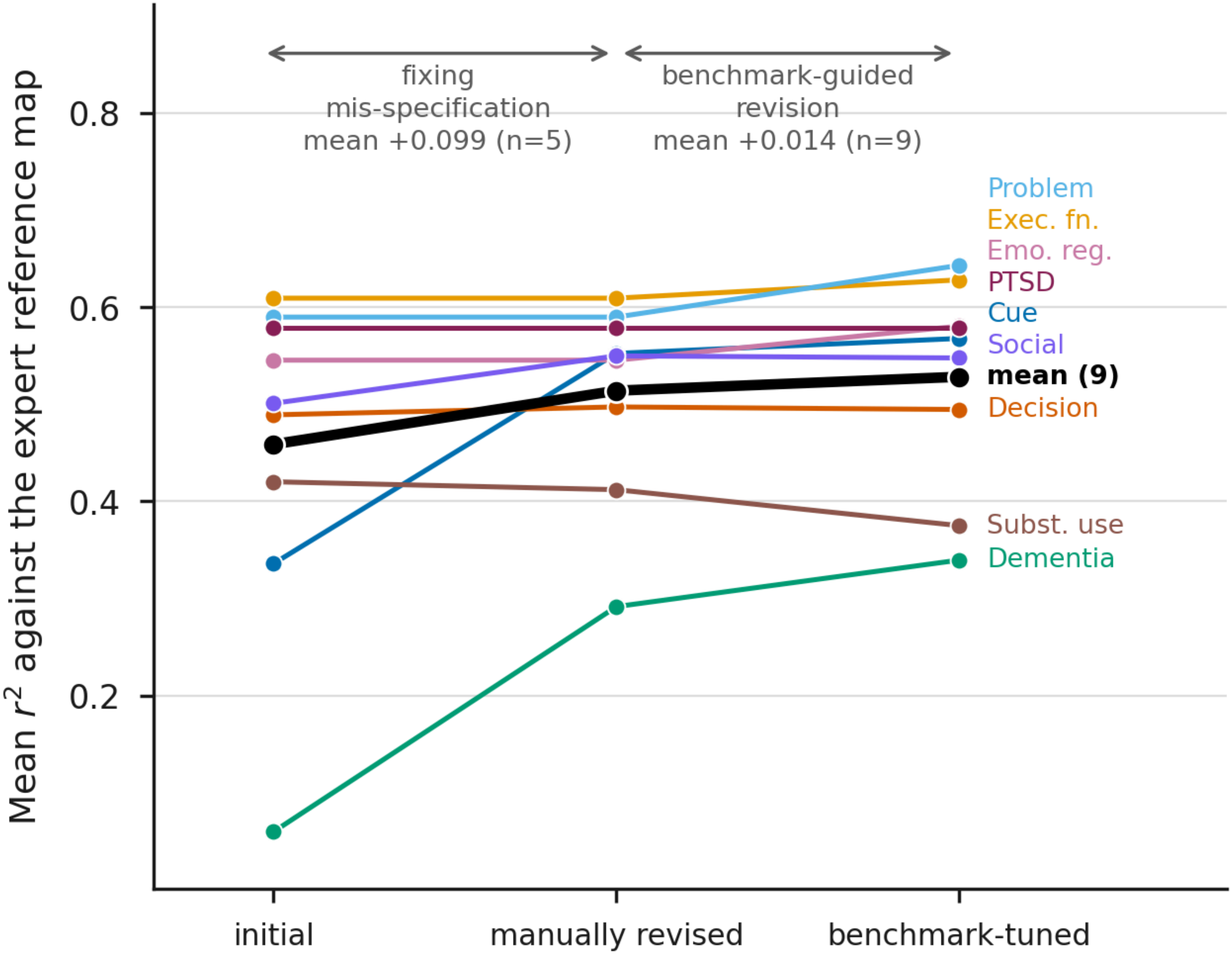

fixing
mis-specification
mean +0.099 (n=5)
benchmark-guided
revision
mean +0.014 (n=9)
Mean $r^2$ against the expert reference map
0.8
0.6
0.4
0.2
Problem
Exec. fn.
Emo. reg.
PTSD
Cue
Social
mean (9)
Decision
Subst. use
Dementia
initial
manually revised
benchmark-tuned

**Fig. S1 | Source-based corrections account for most improvement across criteria revisions.** Mean spatial correspondence ($r^2$) between expert maps and maps from initial, manually revised and benchmark-tuned configurations. Initial configurations encoded eligibility criteria from the reference publications. Revised configurations incorporated manual revisions, primarily from re-reading the source publications and inspecting a small set of miscategorised studies; benchmark-tuned configurations incorporated revisions informed by source publications and benchmark error reports, with LLM assistance in eight of nine review articles. Mean improvement was +0.099 across five pre-tuning revisions and +0.014 across nine tuning transitions, including four review articles tuned directly from their initial configuration. Post hoc classification linked all five pre-tuning revisions (two mis-transcribed criteria, one omitted criterion, one ambiguous criterion, one search-query change) and six of nine tuning revisions (three omitted criteria, two criteria contradicted by the source's own included studies, one ambiguous criterion) to source-based corrections or search changes. The remaining three tuning revisions lacked a direct source basis and occurred only in the benchmark-tuned criteria. Across eight review articles with paired initial and best configurations, mean improvement was +0.077, with increases in seven review articles. The best-performing configuration used in the principal evaluation was selected from the three tiers and is not plotted separately.

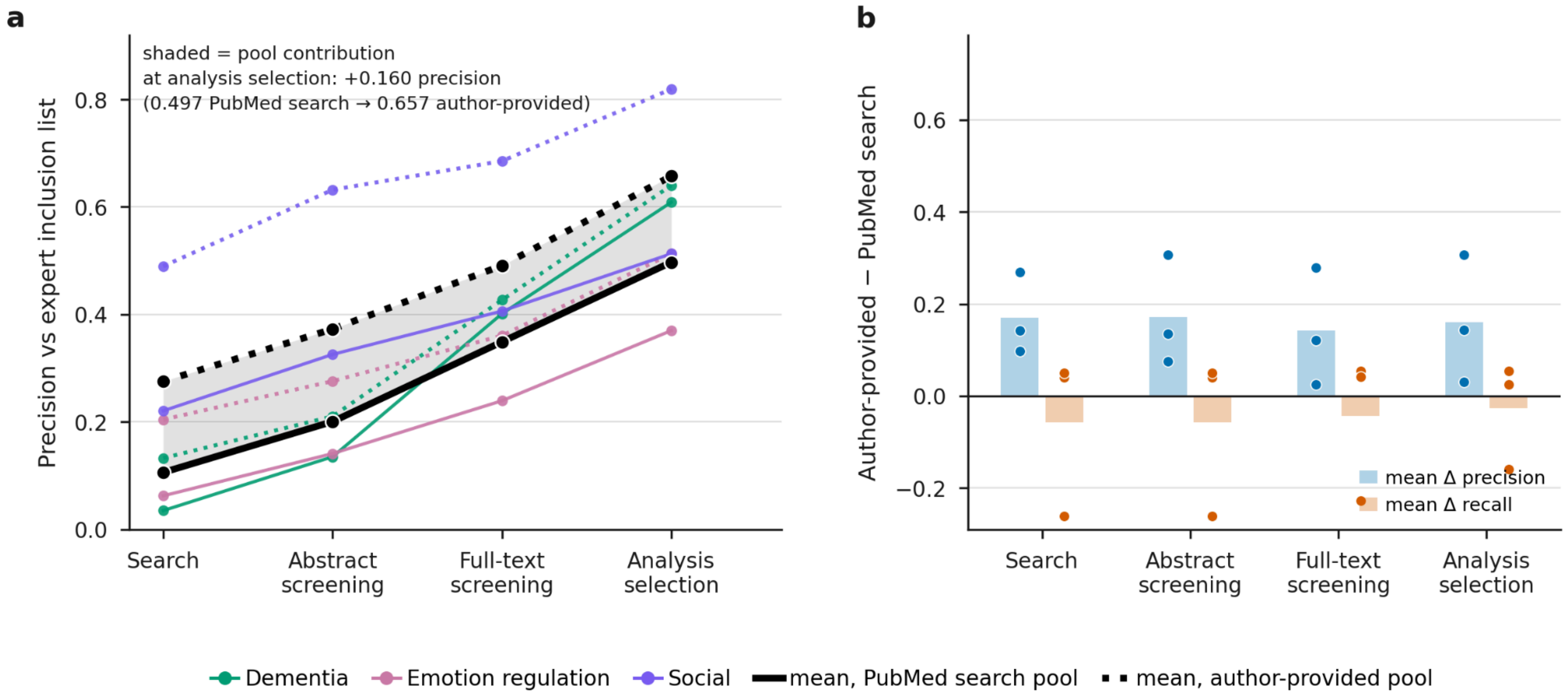


**Fig. S2 | Candidate article pools influence measured screening precision. a, Precision against expert inclusion lists** when the same eligibility criteria are applied to articles retrieved by AutoNIMA's PubMed searches (solid lines) or candidate screening lists obtained from the original reviewers (dotted lines). Author-provided lists were available for three review articles: dementia, emotion regulation, and social cognition. At the final stage, articles were retained if at least one analysis was selected for a meta-analysis. Mean final precision increased from 49.7% to 65.7% when author-provided lists were used, a difference of 16 percentage points. This comparison indicates that differences between the candidate pools considered by automated and original reviewers can lower apparent precision against published inclusion lists. b, Differences in precision and recall at each stage between the reviewer-provided candidate pool and the PubMed search pool. Points show differences for individual review articles, and bars show the mean across the three review articles. Recall decreased in one review article because the reviewer-provided list covered only a subset of the originally screened articles.

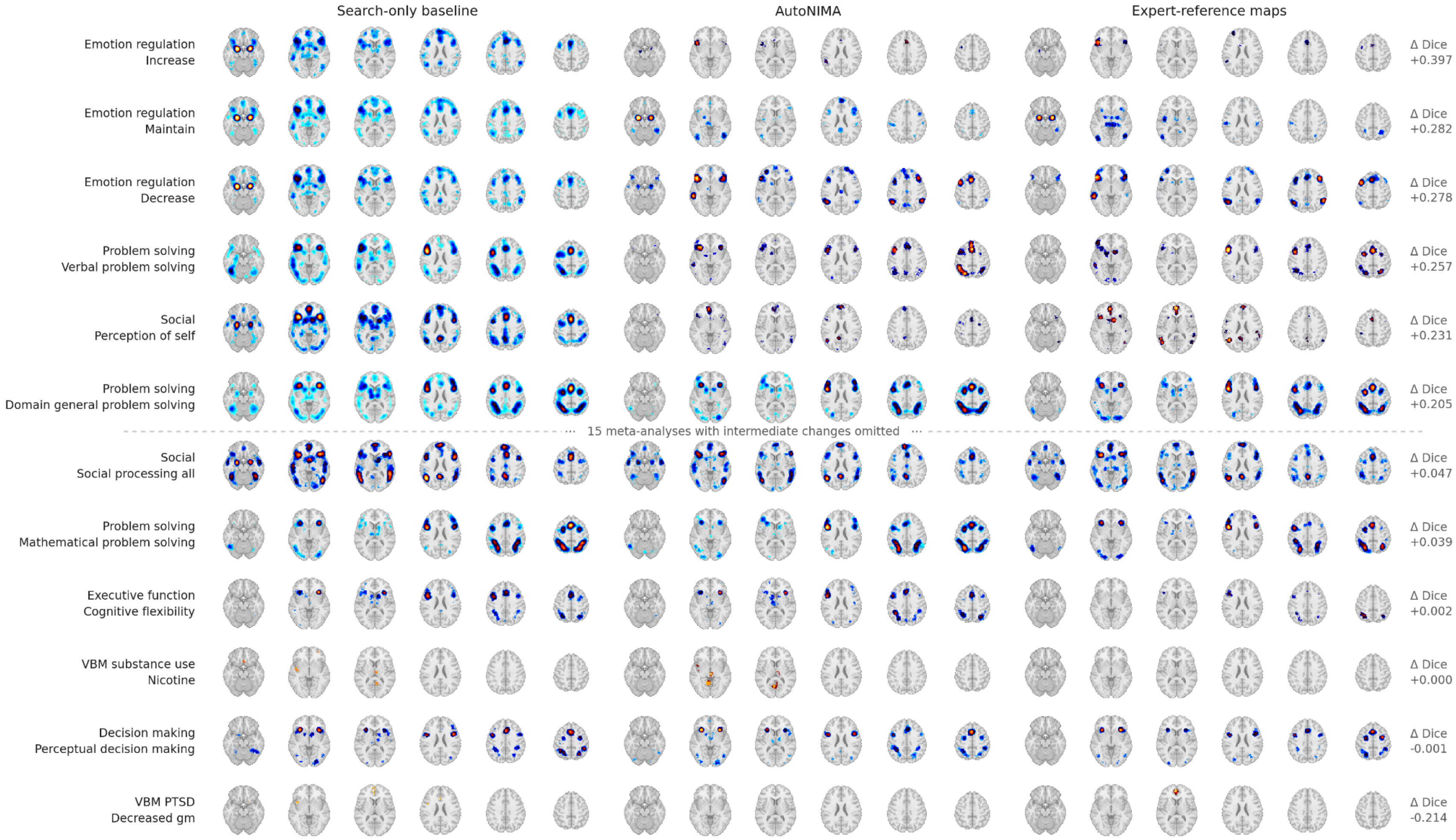


**Fig. S3 | Search-only baseline, AutoNIMA, and expert-reference maps for the most and least improved meta-analyses.** Axial slices at four levels show, from left to right, the search-only baseline, AutoNIMA (labelled "Full pipeline") and expert-reference maps for the top and bottom 6 meta-analyses included in Fig. 4. Meta-analyses are sorted from greatest to smallest improvement in Dice overlap with the expert-reference map for AutoNIMA relative to the search-only baseline. Maps are displayed at $z > 1.96$, the threshold used for the reported overlap analyses. Squared voxelwise correlations were calculated from unthresholded maps.

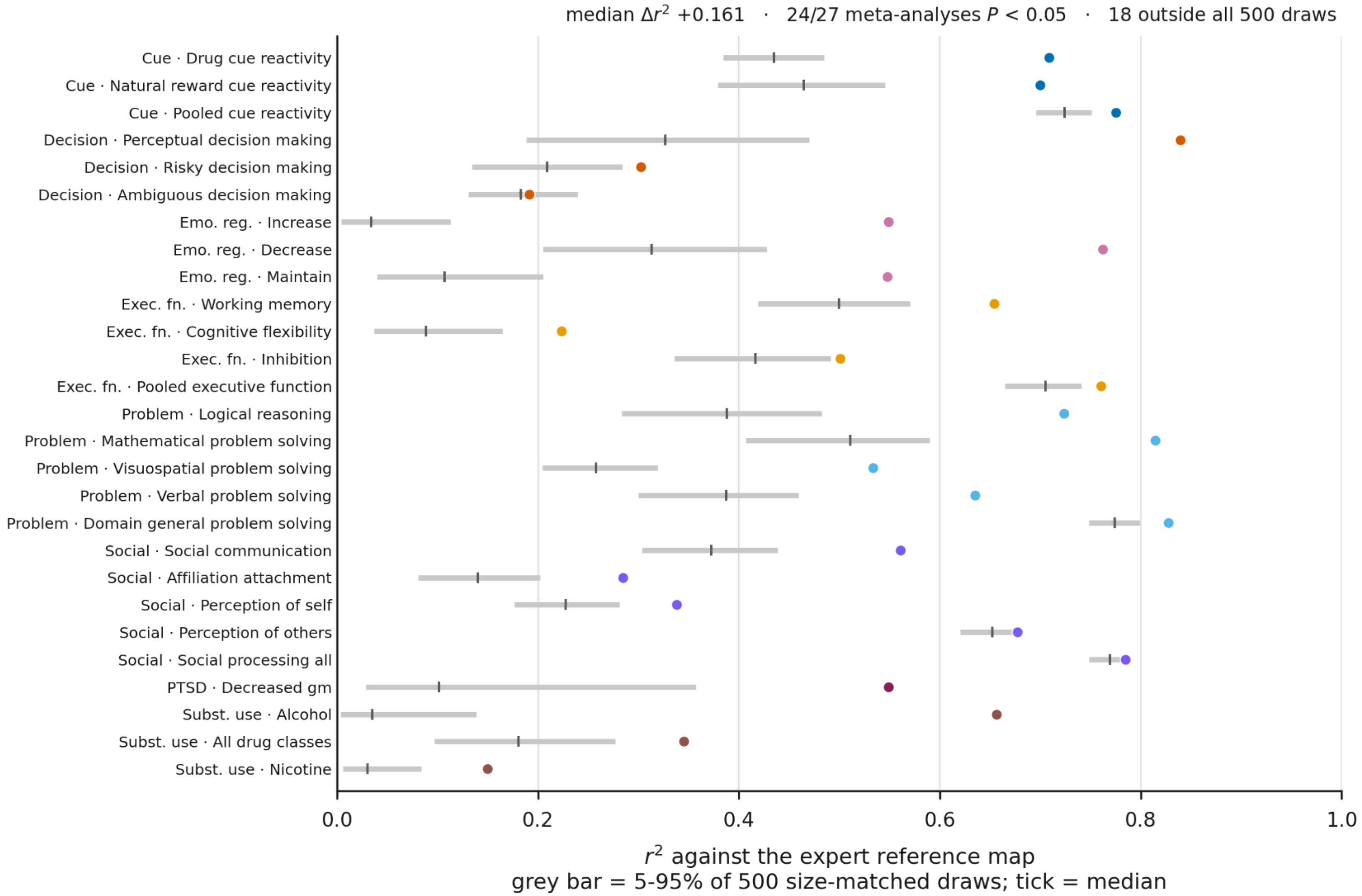

median Δ$r^2$ +0.161 · 24/27 meta-analyses $P < 0.05$ · 18 outside all 500 draws
Cue · Drug cue reactivity
Cue · Natural reward cue reactivity
Cue · Pooled cue reactivity
Decision · Perceptual decision making
Decision · Risky decision making
Decision · Ambiguous decision making
Emo. reg. · Increase
Emo. reg. · Decrease
Emo. reg. · Maintain
Exec. fn. · Working memory
Exec. fn. · Cognitive flexibility
Exec. fn. · Inhibition
Exec. fn. · Pooled executive function
Problem · Logical reasoning
Problem · Mathematical problem solving
Problem · Visuospatial problem solving
Problem · Verbal problem solving
Problem · Domain general problem solving
Social · Social communication
Social · Affiliation attachment
Social · Perception of self
Social · Perception of others
Social · Social processing all
PTSD · Decreased gm
Subst. use · Alcohol
Subst. use · All drug classes
Subst. use · Nicotine
0.0
0.2
0.4
0.6
0.8
1.0
$r^2$ against the expert reference map
grey bar = 5-95% of 500 size-matched draws; tick = median

**Fig. S4 | Analysis-level selection improves map recovery relative to equally sized random subsets.** Observed correspondence with expert-reference maps ($r^2$) compared with a random-selection distribution for each of 27 meta-analyses. Gray bars span the 5th–95th percentiles. For each meta-analysis, 500 subsets were sampled from the same candidate analysis pool, each containing the same number of analyses as the selected set. MKDA was repeated for every subset using the same settings as the observed synthesis. This comparison controls the number of pooled analyses while varying which analyses are included. Across the 27 meta-analyses, the median difference between the observed $r^2$ and the mean $r^2$ of the corresponding random subsets was +0.161. Ticks mark the median of each random-subset distribution. Observed correspondence was significantly higher than expected under random selection in 24 of 27 meta-analyses, both before and after Benjamini–Hochberg correction (uncorrected $P < 0.05$, corrected $q < 0.05$). Empirical $P$ values were calculated as one plus the number of random subsets achieving at least the observed correspondence, divided by 501; in 18 meta-analyses no random subset matched the observed result, giving the minimum attainable value of 0.002. Percentile ranges describe variation across random subsets and are not confidence intervals for the observed result.

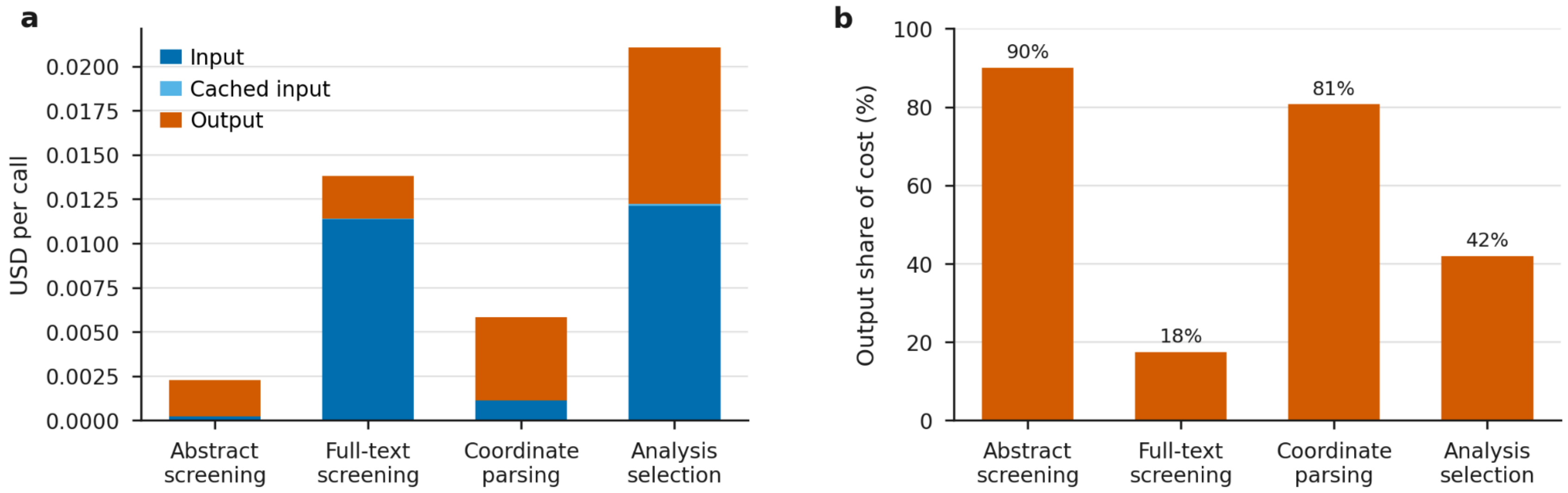


**Fig. S5 | Model-use costs estimated from recorded token consumption.** a, Mean cost per LLM call at each workflow stage, broken down by uncached input, cached input and output token charges. b, Percentage of per-call costs attributable to output tokens at each stage. Analysis selection is labelled "Annotation" in both panels. Costs were calculated from recorded token consumption and the applicable token prices (Methods). Mean per-call costs were US$0.0138 for full-text screening and US$0.0211 for analysis selection. Output tokens accounted for most of the abstract-screening cost. Applying measured usage rates to the observed number of operations at each stage gave an estimated total cost of US$194 across all nine review articles, approximately US$21.6 per review article. The median cost per contributing study across review articles was US$0.085. Totals for each review article account for differences in the number of articles reaching each stage. These estimates cover model use and exclude article-access charges, other computing costs and researcher time.

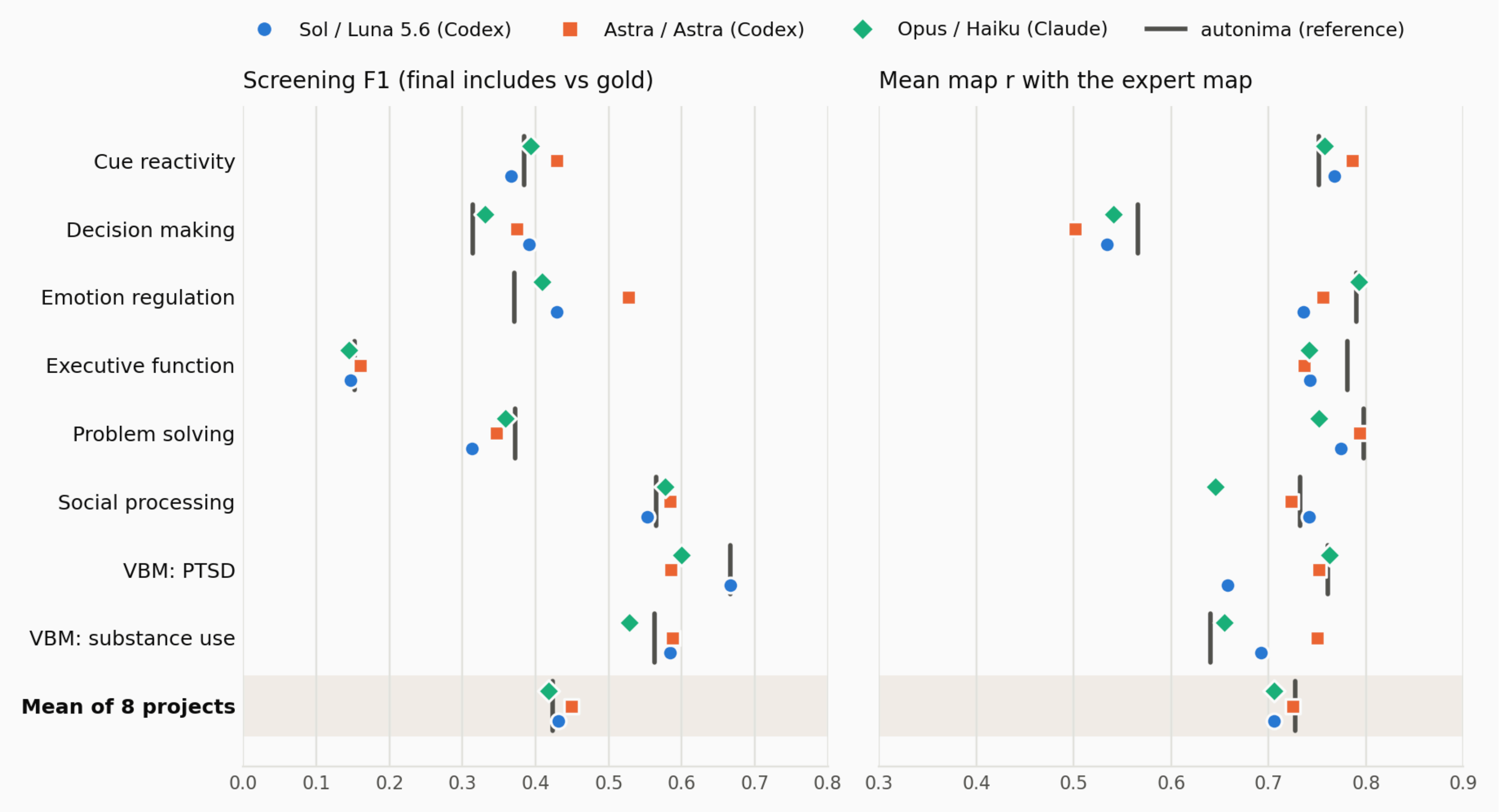


**Fig. S6 | Agreement of three skills-based workflows with the expert meta-analyses, by review article. Each row shows one of the eight review articles included in the spatial comparison. The shaded row shows the mean across the eight displayed review articles.** Left: screening F1 of each workflow's final included studies against the expert-curated inclusions. Right: mean Pearson correlation ($r$) between each workflow's unthresholded MKDA $z$ map and the expert-reference map, averaged over the review article's eligible meta-analyses. This comparison used the original evaluation set (Methods). Markers show the three workflows, labelled orchestrator / judge model: Sol / Luna 5.6 and Astra / Astra (Codex), and Opus / Haiku (Claude Code). The grey tick marks the AutoNIMA pipeline as a reference.
